# Single-cell RNA-seq reveals endoimmune cells in zebrafish

**DOI:** 10.1101/2019.12.31.892240

**Authors:** Yunwei Shi, Peipei Qian, Jiajing Sheng, Xu Zhang, Xiaoning Wang, Jingxiang Zhao, Guanyun Wei, Xia Liu, Gangcai Xie, Dong Liu

**Affiliations:** School of life science, Key Laboratory of Neuroregeneration of Jiangsu and Ministry of Education, Co-innovation Center of Neuroregeneration, Nantong University, Nantong, China; Medical school, Nantong University, Nantong, China

**Keywords:** endoimmune cells, endothelial cells, single-cell RNA sequencing, zebrafish, inflammation

## Abstract

Endothelial cells (ECs) constitute a monolayer that covers the interior surface of blood vessels and participates in various processes. Although vascular ECs share certain common properties, they differ in both structure and function. So far, the transcriptome profile and heterogeneity of the full repertoire of ECs in vertebrates remain poorly understood. The relatively small size of zebrafish embryos and larvae allows a feasible analysis of the broad spectrum of ECs within every tissue and organ of a whole organism. ECs have been suggested to be conditional innate immune cells. Whether ECs possess the comparable capacity of involvement in immune response is so far undetermined. Currently, through single-cell RNA sequencing analysis of total ECs of zebrafish we identified a fraction of endothelial cells expressing the marker genes of innate immune cells, named “endoimmune cells”. We found the percentage of these cells gradually increased along with the embryonic development. Then, we observed the patrolling mCherry+ cells displayed the morphology alike to the macrophages and neutrophils. Furthermore, we revealed that some of the kdrl:ras-mCherry ECs were labelled with coro1a:EGFP as well. In addition, we demonstrated that the mCherry+ EC from intersegmental vessel could gradually present the expression of GFP in *Tg(kdrl:ras-mCherry∷coro1a:GFP)* line, suggesting the endoimmune cells are derived from ECs. Importantly, we showed the endoimmune cells are responsive to the inflammation in zebrafish. Taken together, these data suggested the existence of endoimmune cells, a novel type of subpopulations of ECs. It will provide novel insights for understanding endothelial roles in both normal physiological function and human diseases and enable endoimmune cells-based target therapies.

---

Evolutionary emerging of the cardiovascular system endowed a means of aiding the diffusion process allowing the development of larger organisms [1]. The blood vessels serve to provide rapid transport of nutrients around the body and removal of waste products for animals and human beings. They disseminate in all organs of the body and play a variety of roles in embryonic development and medical conditions [2]. Endothelial cells (ECs) form a subtle monolayer that covers the interior surface of all blood vessels and participates in a large number of physiological processes [3]. Endothelial dysfunction is implicated in pathogenesis of various diseases, including atherogenesis, hypertension, diabetes mellitus, and vascular inflammation [4, 5]. Although vascular ECs share certain common properties, they differ in both structure and function due to the biomechanical or biochemical environmental cues [6]. Characterizing functional subpopulations of ECs will provide novel insights for understanding endothelial roles in both of normal physiological function and human diseases with vascular dysfunction and enable sub-population ECs based target therapies. So far, the transcriptome profile and heterogeneity of the full repertoire of ECs in vertebrates remain poorly understood. The relatively small size of zebrafish embryos and larvae allows feasible analysis of the broad spectrum of ECs from every tissue and organ in a whole organism.

For the first time, we carried out the single-cell RNA sequencing (scRNA-seq) analysis of total ECs from zebrafish at 4 developmental stages, including 24 hpf (hours post fertilization), 48 hpf, 4 dpf (days post fertilization) and 8 dpf. Since endothelial cells represent a small fraction of zebrafish embryos, it makes analysis of these cells difficult. Therefore, we employed transgenic zebrafish *Tg(kdrl:EGFP)* (Figure S1), in which EGFP was expressed under control of the *kdrl* promoter in ECs [7]. Following proteolytic dissociation of embryos, fluorescence activated cell sorting (FACS) isolated GFP+ cell populations. For each stage, around 300-500 zebrafish embryos or larvae were used for the ECs collection. The purity of these sorted cells was validated by diagnostic FAC resorting and Taqman PCR analysis of marker genes of the ECs and confirmed by microscopic inspection. The isolated ECs were analyzed using Large-scale scRNA-seq (10× Genomics) platform (Figure 1a). Several criteria were applied to select the single cells, including only keeping the genes that are expressed (Unique Molecular Identifiers or UMI larger than 0) at least in 3 single cells, selecting single cells with the number of expressed genes at the range between 500 and 3000, and requiring the percentage of sequencing reads on mitochondrial genome being less than 5 percentage (Figure S2). A total of 14168 ECs was captured, with each stage over around 2000 ECs (Figure S3). Through clustering analysis of gene expression, we found that these ECs were categorized into 16 subpopulations (Figure 1b; Figure S4). The *vascular endothelial growth factor receptor 2* (*kdrl*) displayed heterogeneous expression by different cluster (Figure S5). The cluster one showed the highest expression of *kdrl* and the enrichment of some other canonical EC markers *dll4*, *egfl7*, and *flt1* etc. Interestingly, we found that cluster 10 and 11 were defined by higher expression of many innate immune cells’ markers, including *lyz*, *mpx*, *coro1a*, *mfap4*, and *marco* etc. (Figure 2). To investigate the cellular functions of the two EC subpopulations, we carried out the specific genes’ enrichment analysis for each of the two subpopulations. Several immune relevant processes and activities were enriched in these two clusters, including response to bacterium, immunity, immune system process and innate immunity in the cluster 10 (Figure 3a), while inflammatory response, response to bacterium, immune response and macrophage chemotaxis in the cluster 11 (Figure 3b), suggesting these two EC subpopulations possess innate-immunity signatures. Based on these analyses, we named these cells “endoimmune cells”.

**Figure 1:**
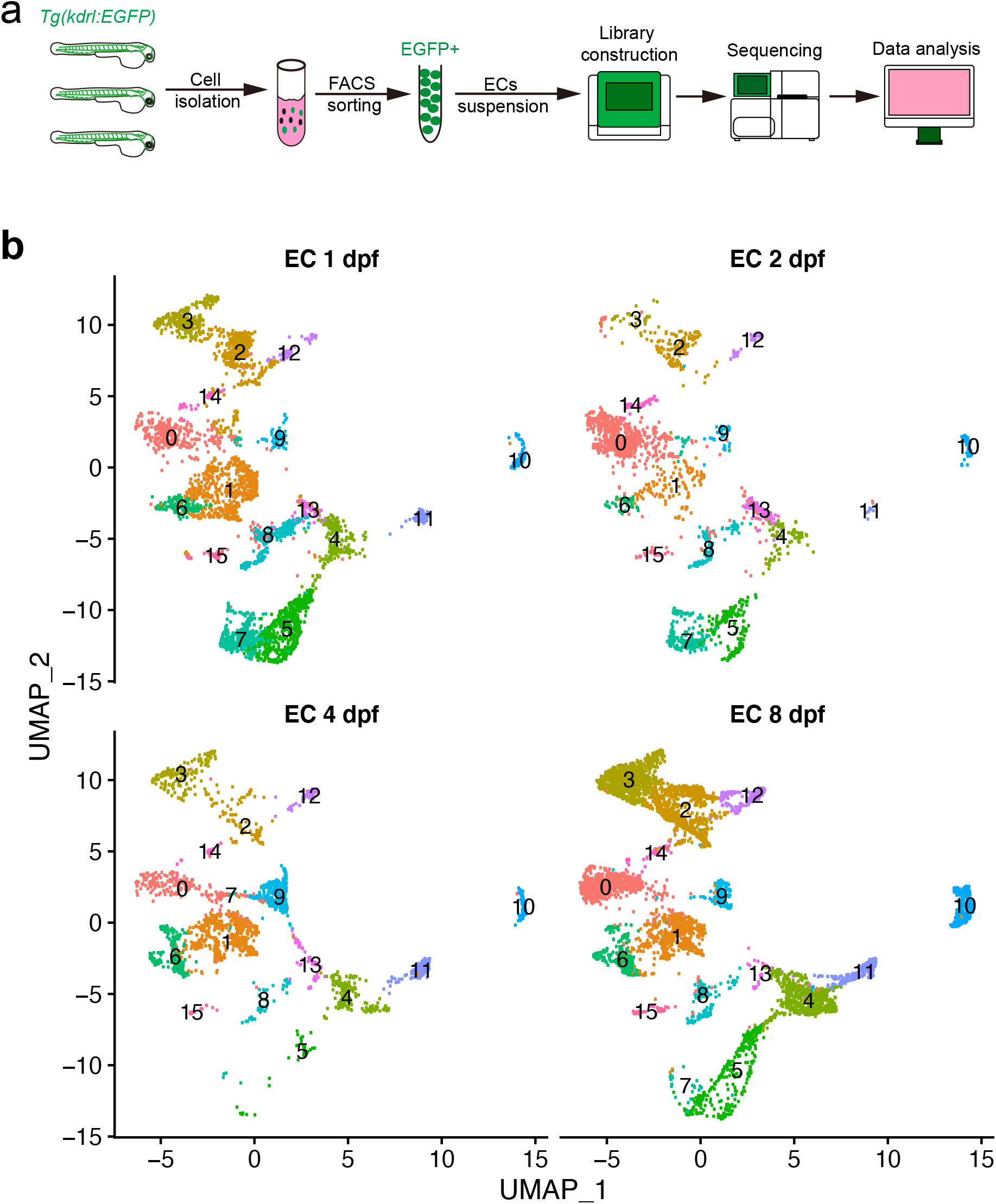
(a) Workflow of single cell RNA sequencing study of zebrafish endothelial cells. (b) UMAP representation of each endothelial cell (EC) subpopulation at four developmental stages. In total, 15 EC subpopulations were found.

**Figure 2:**
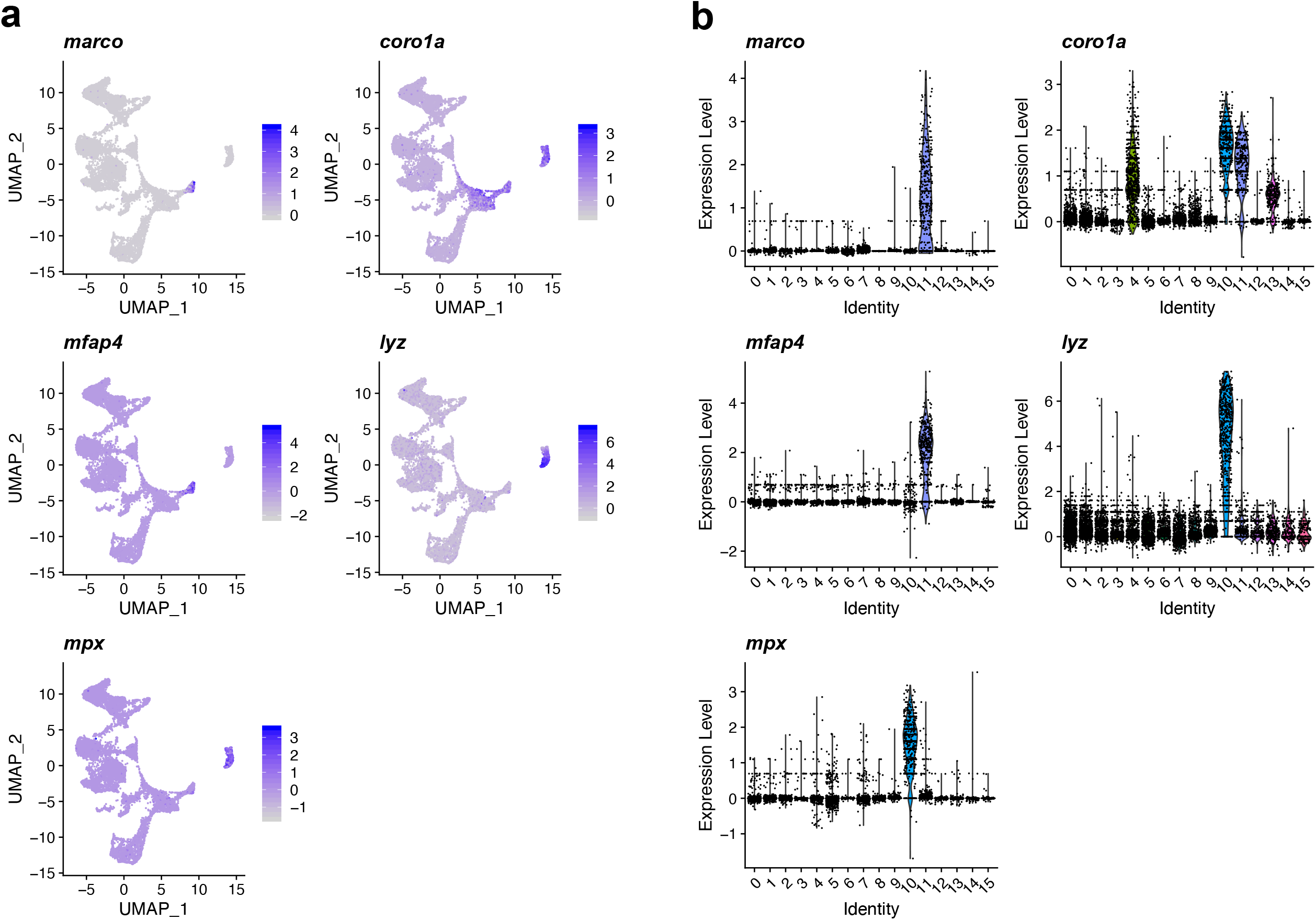
(a) EC subpopulation expression pattern of five innate immune cell marker genes.(b) Violin plot representation of the expression of innate immune cell marker genes in each EC subpopulation. EC subpopulation cluster 10 and cluster 11 were enriched with the innate cell marker gene RNAs.

**Figure 3:**
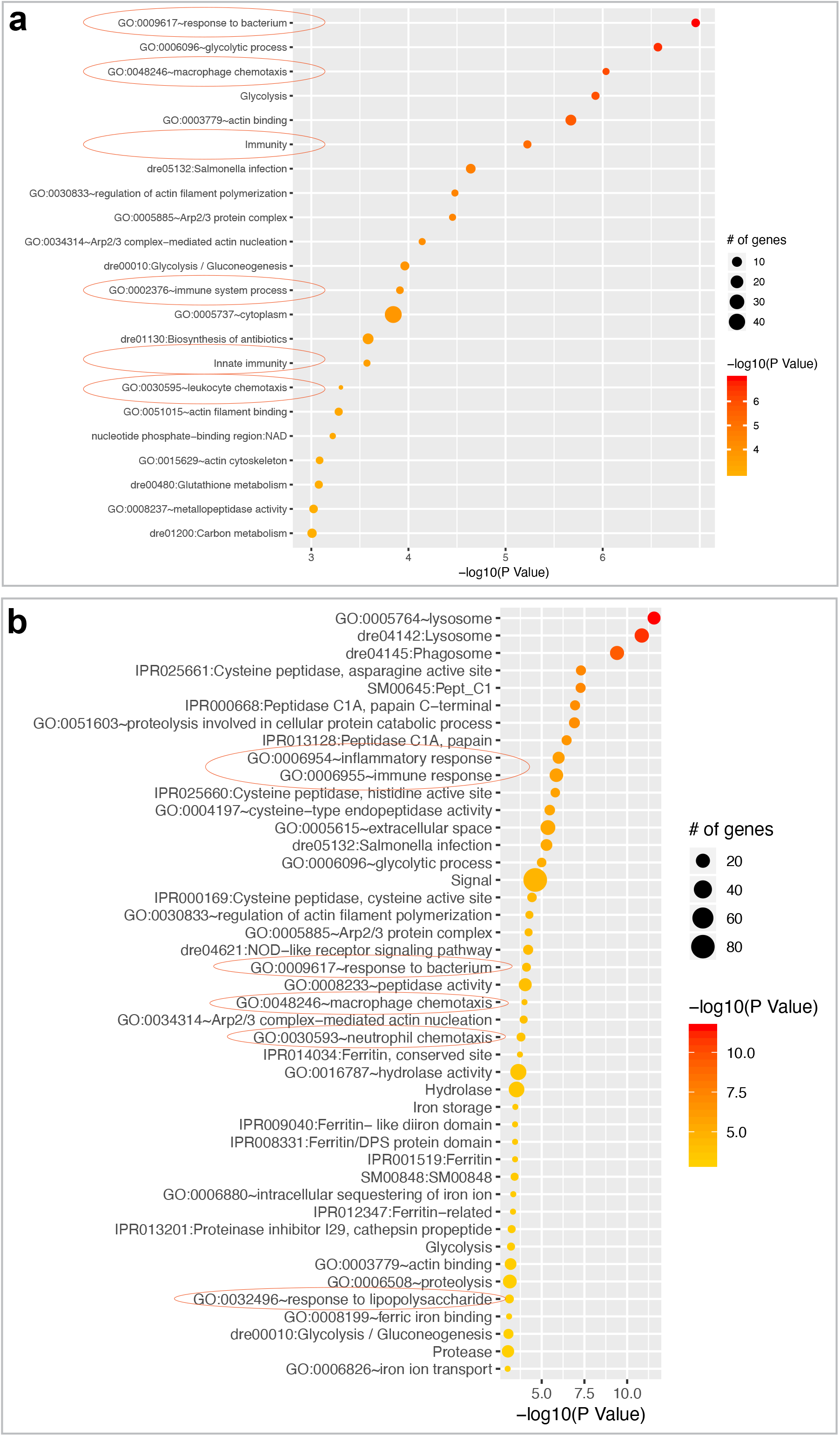
DAVID gene enrichment analysis of endoimmune EC subpopulations, and the dotplot representation of the enrichments for EC subpopulation cluster 10 (a) and cluster 11 (b). “Innate immunity”, “immune response” etc immune-related terms were enriched.

To examine whether these two EC subpopulations are contained in total ECs at different developmental stages, we analyzed the subpopulations at 24 hpf, 48 hpf, 4 dpf and 8 dpf. It was revealed that the percentage of these cells gradually increased along with the embryonic development (Figure 4), supporting that these cells are not the immature endothelial progenitor cells, which potentially are able to differentiate into innate-immunity cells. Furthermore, we did the confocal microscopic live-imaging analysis of *Tg(kdrl:ras-mCherry)* line and observed the patrolling mCherry+ cells displayed the morphology alike to the macrophages and neutrophils in the embryos at 48 hpf and 72 hpf stages (Figure S6). Moreover, we observed that double-labelled cells by both mCherry and GFP in *Tg(kdrl:ras-mCherry∷coro1a:GFP)* line (Figure 5). These results suggest the existence of endoimmune cells, in which both endothelial marker genes and innate-immune cell marker genes were detected. In addition, the confocal microscopic live-imaging analysis of *Tg(kdrl:ras-mCherry∷coro1a:GFP)* line showed that the mCherry+ EC from intersegmental vessel could gradually present the expression of GFP (Figure 6, Movie 1), suggesting the endoimmune cells are derived from ECs.

**Figure 4:**
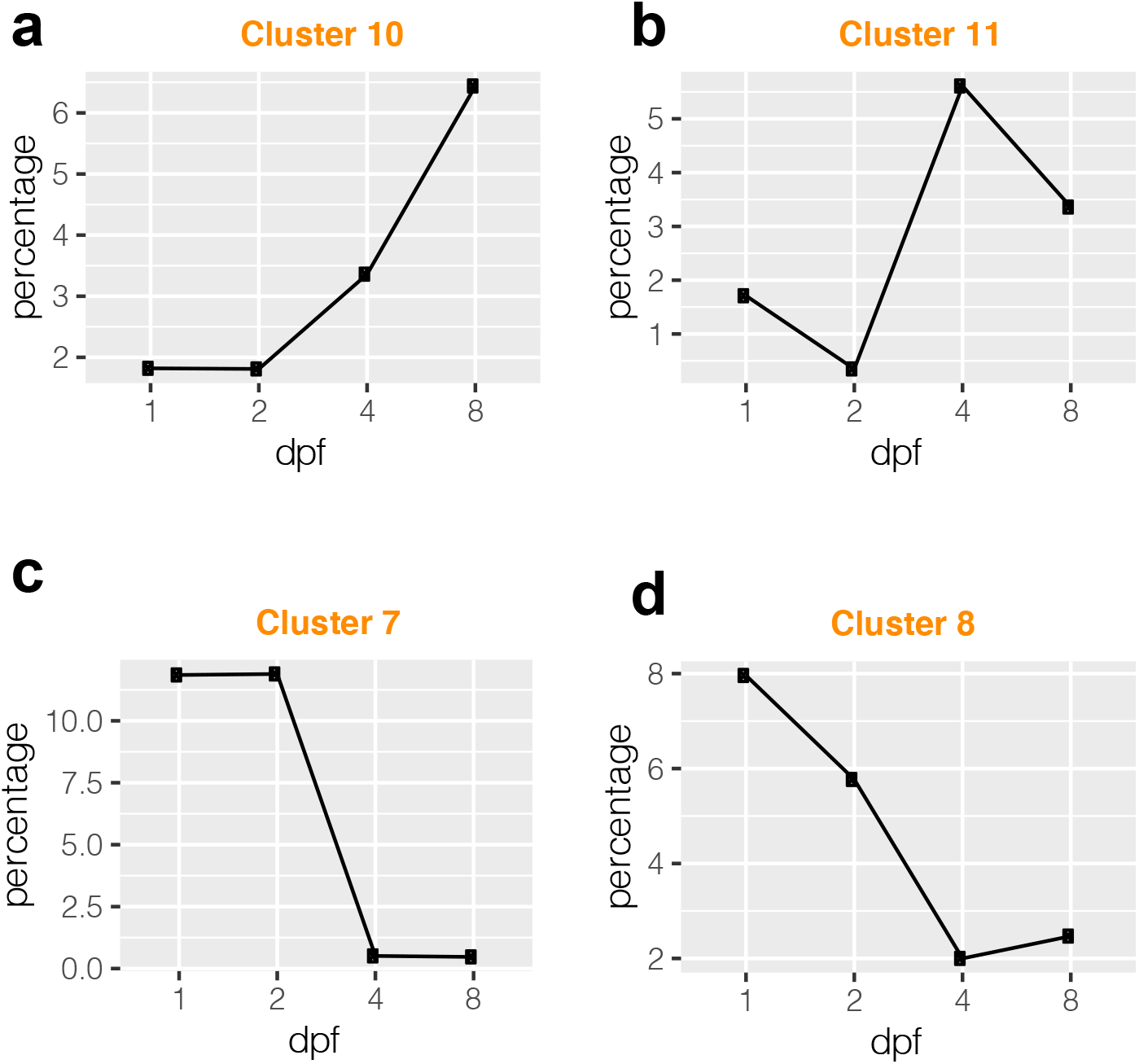
Developmental changes of the cell proportions for the four EC subpopulations. Increasing of EC cluster 10 (a) and cluster 11 (b) cell populations at 4dpf and 8dpf comparing to the earlier developmental time. Decreasing of EC cluster 7 (c) and cluster 8 (d) cell populations at 4dpf and 8dpf comparing to the earlier developmental time. EC cluster 10 and cluster 11 are endoimmune cells.

**Figure 5:**
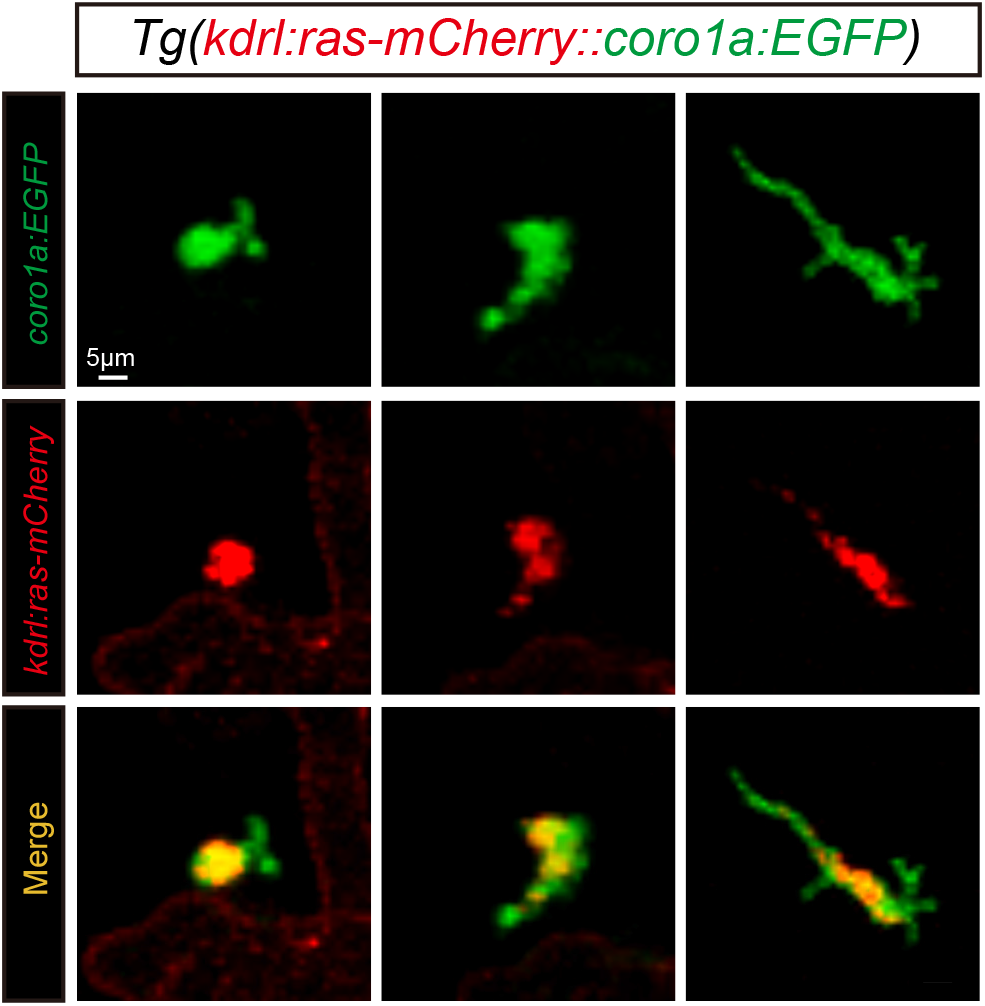
Confocal imaging analysis of mCherry and EGFP double positive cells in *Tg(kdrl:ras-mCherry∷coro1a:EGFP)* line.

**Figure 6:**
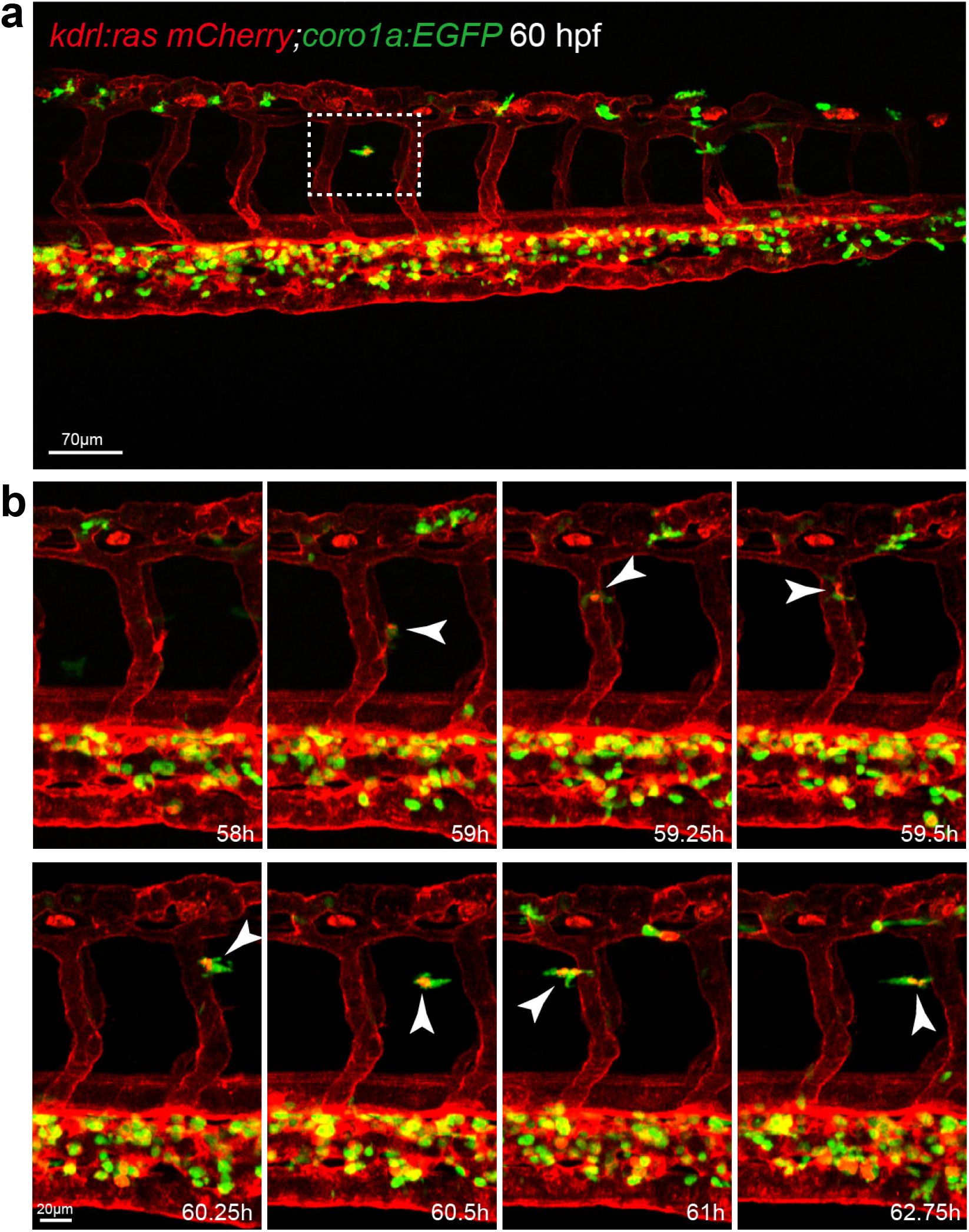
Confocal time-lapse imaging analysis of emerging expression of EGFP in mCherry positive cells in *Tg(kdrl:ras-mCherry∷coro1a:EGFP)* line.

As endoimmune cells were characterized with the enrichment of the genes relevant to innate immunity and immune response, we reasoned that these cells are involved in the inflammatory response in zebrafish. To determine whether this was the case, we examined the presence of endoimmune cells in response to the inflammation. We microinjected Lipopolysaccharides (LPS) and Zymosan (ZYM), which are used to cause inflammatory reaction, respectively into 48 hpf *Tg(kdrl:ras-mCherry∷coro1a:GFP)* embryo and did the confocal microscopic imaging analysis. We found that the microinjection of both LPS and ZYM efficiently stimulated the accumulation of macrophage and neutrophils in the injected location (Figure S7), indicating the activation of innate immune system. Importantly, the endoimmune cells were observed in the injection sites as well (Figure 7), suggesting that these cells are responsive to the inflammation in zebrafish.

**Figure 7:**
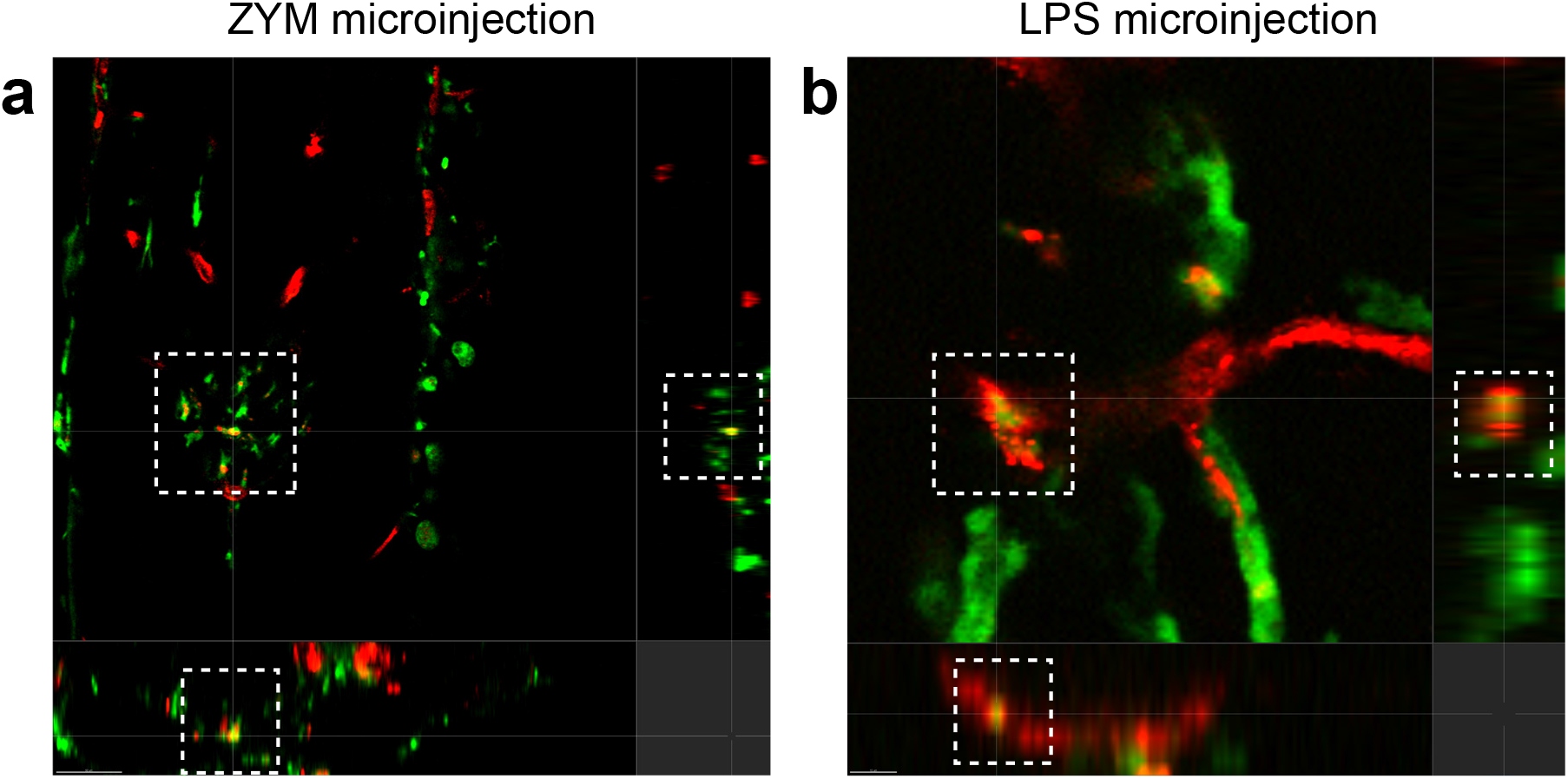
Confocal imaging analysis of mCherry and EGFP double positive cells in ZYM microinjected (a) and LPS microinjected (b) *Tg(kdrl:ras-mCherry∷coro1a:EGFP)* line.

ECs have been suggested to be conditional innate immune cells, which actively participate in both innate and adaptive immune responses [8]. ECs at a site of inflammation could function as both active immune effectors and regulators of inflammatory processes [9]. It was reported that bacterial endotoxin lipopolysaccharide (LPS) treatment caused the endothelial activation, defined as the acquisition of new capacities by resting endothelial cells. [9–12]. In addition, the pro-inflammatory cytokines and chemokines secreted by the stimulated ECs played important roles in provoking the immune response by recruiting immune cells to the site of inflammation [8, 9, 13]. Whether total ECs possess the comparable capacity of involvement in immune response is so far undetermined. Currently, we identified a fraction of endothelial cells expressing the marker genes of innate immune cells, named endoimmune cells in zebrafish. Thus, we suppose the endoimmune cells might play more important roles in response to inflammation than the rest of the ECs. The discovery of endoimmune cells contributes to enlarge the knowledge of spectrum of endothelial heterogeneity and will provide a novel insight to understand the function of ECs in inflammatory response and infectious diseases. Furthermore, it might enable endoimmune cells-based target therapies.

## Materials and methods

### Zebrafish

All zebrafish experimentation was carried out in accordance with the NIH Guidelines for the care and use of laboratory animals (https://oacu.oir.nih.gov/regulations-standards). In addition, the study was conducted conforming to the local institutional rules and the Chinese law for the Protection of Animals. Zebrafish embryos were obtained through natural mating (AB line) and maintained under standard conditions as we previously described [14]. Embryonic stages were defined as our previous work [15]. Embryos for imaging analysis after 24 hpf were treated with 0.2mM 1-phenyl-2-thio-urea (PTU, a tyrosinase inhibitor commonly used to block pigmentation and aid visualization of zebrafish development Sigma, P7629). Transgenic zebrafish lines, *Tg(kdrl:EGFP)*, *Tg(kdrl:ras-mCherry)* and *Tg(coro1a:EGFP)* were used as described in previous work [14, 16, 17].

### Fluorescence-activated cell sorting (FACS) and Single-cell RNA sequencing

*Tg(kdrl:EGFP)* embryos were raised in E3 solution to the indicated developmental stage and dechorionated by forceps. Embryos were transferred into a 15 mL or 50 mL falcon tube with 10 mL or 25 mL phosphate buffered saline (PBS) containing 0.25% trypsin and incubated in a rolling machine for 30 min at 28°C during which they were pipetted up and down per 5 min.

After centrifuging for 5 min at 800 g at 4°C, cells were resuspended in PBS containing 0.25% trypsin and 15% fetal calf serum (FCS) to stop the digestion and centrifuged for 5 min at 800 g at 4°C. Cells were rinsed with PBS containing 2% FCS for 2 times and resuspended in PBS. FACS was performed on a FACS Aria3 (BD Biosciences). Kdrl:EGFP positive cells were first enriched by fluorescence-activated cell sorting (FACS), and then went through library preparation following the standard protocol named Chromium Single Cell 3’Reagent Kits v3 from 10X Genomics Company. The 10× Chromium single cell libraries were sequenced by Illumina HiSeq4000 sequencer.

### Imaging

For confocal live imaging analysis of zebrafish embryos, they were anaesthetized with E3 solution/0.16 mg/mL tricaine/1% 1-phenyl-2-thiourea (Sigma) and embedded in 0.5-0.8% low melting agarose. Confocal live imaging analysis was performed with a Nikon TI2-E-A1RHD25 microscope. Analysis was performed using Imaris software.

### Lipopolysaccharides (LPS) and Zymosan (ZYM) microinjection

Around 2 nL LPS (Sigma, L5024, 1 mg/mL), ZYM (Sigma, Z4250, 10 mg/ml) and PBS used as a control were microinjected into the somites of zebrafish embryos at 48 hpf or other indicated stages.

### Single-cell gene expression profile analysis

Cell Ranger 3.0.2 (https://github.com/10XGenomics/cellranger) was used to convert the raw sequencing data to single-cell level gene counts matrix. The clustering of single cells and the marker genes in each cluster were analyzed by Seurat 3.0 (https://satijalab.org/seurat/install.html) [18]. Several criteria were applied to select the single cells, including only keeping the genes that are expressed (Unique Molecular Identifiers or UMI larger than 0) at least in 3 single cells, selecting single cells with the number of expressed genes at the range between 500 and 3000, and requiring the percentage of sequencing reads on mitochondrial genome being less than 5 percentage. Furthermore, sctransform method [19] were applied to remove technical variation, and ClusterProfiler [20] was used to do the Gene Ontology enrichment analysis based on the marker genes of each cell cluster. The detail information about the data processing can be found in the source code of this project (https://github.com/gangcai/ZebEndoimmune).

## Supporting information

Movi 1

## Acknowledgements

This study was supported by grants from the National Natural Science Foundation of China (81870359, 2018YFA0801004, 81570447 to Dong Liu; 31900484 to Gangcai Xie; 81970432 to Yunwei Shi), and Natural Science Foundation of Jiangsu Province (BK20180048, 17KJA180008 and BRA2019278 to Dong Liu, BK20190924 to Gangcai Xie).

## Supplementary Figure Legends

**Figure S-1: Confocal imaging analysis of *Tg(kdrl: EGFP)* line at 24 hpf, 48 hpf, 4 dpf, and 9 dpf.**

**Figure S-2: Overview of the number of genes, total UMIs and percentage of mitochondrial UMIs for the single cell RNA sequencing. (a) Before filtering. (b) After filtering. Cell selection criteria: 500< number of genes < 3000; 0< percentage of mitochondrial UMIs < 5%.**

**Figure S-3: Number of single cells used in this study for each developmental stages.**

**Figure S-4: UMAP representation of EC subpopulations. All single cells (after filtering) from the four developmental stages were included in this illustration.**

**Figure S-5: Wide expression of kdrl at all EC subpopulations.**

**Figure S-6: Confocal imaging analysis of the patrolling cells in *Tg(kdrl:ras-EGFP)* line and *Tg(coro1a:EGFP)* line.**

**Figure S-7: Confocal imaging analysis of mCherry and EGFP double positive cells in ZYM microinjected (a) and LPS microinjected (b) *Tg(kdrl:ras-mCherry∷coro1a:EGFP)* line.**

