## Supplementary Figures 1-7 for "Single-cell RNA-seq reveals endoimmune cells in zebrafish"

### Supplementary Figure S1

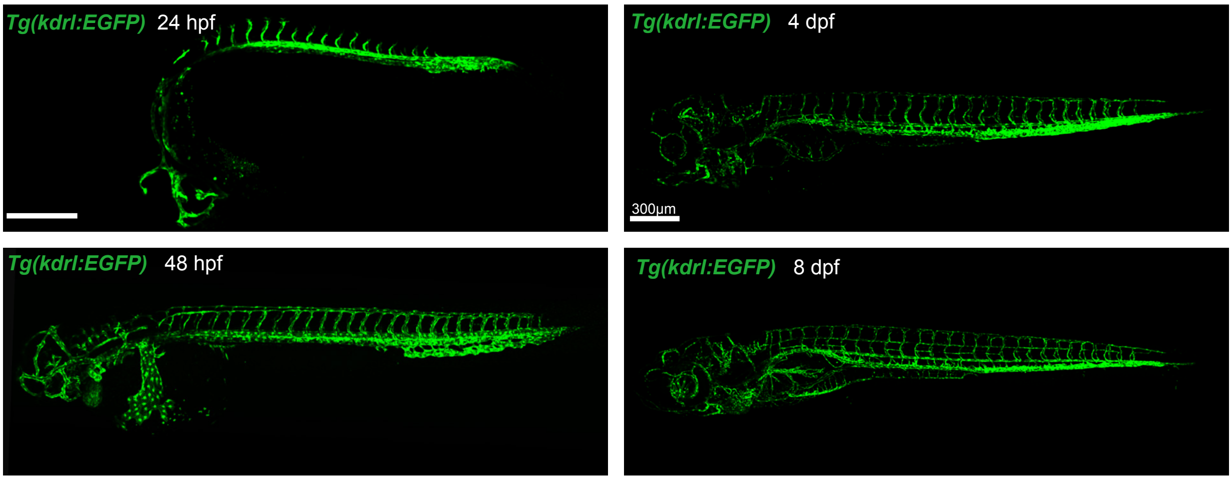

Supplementary Figure S2

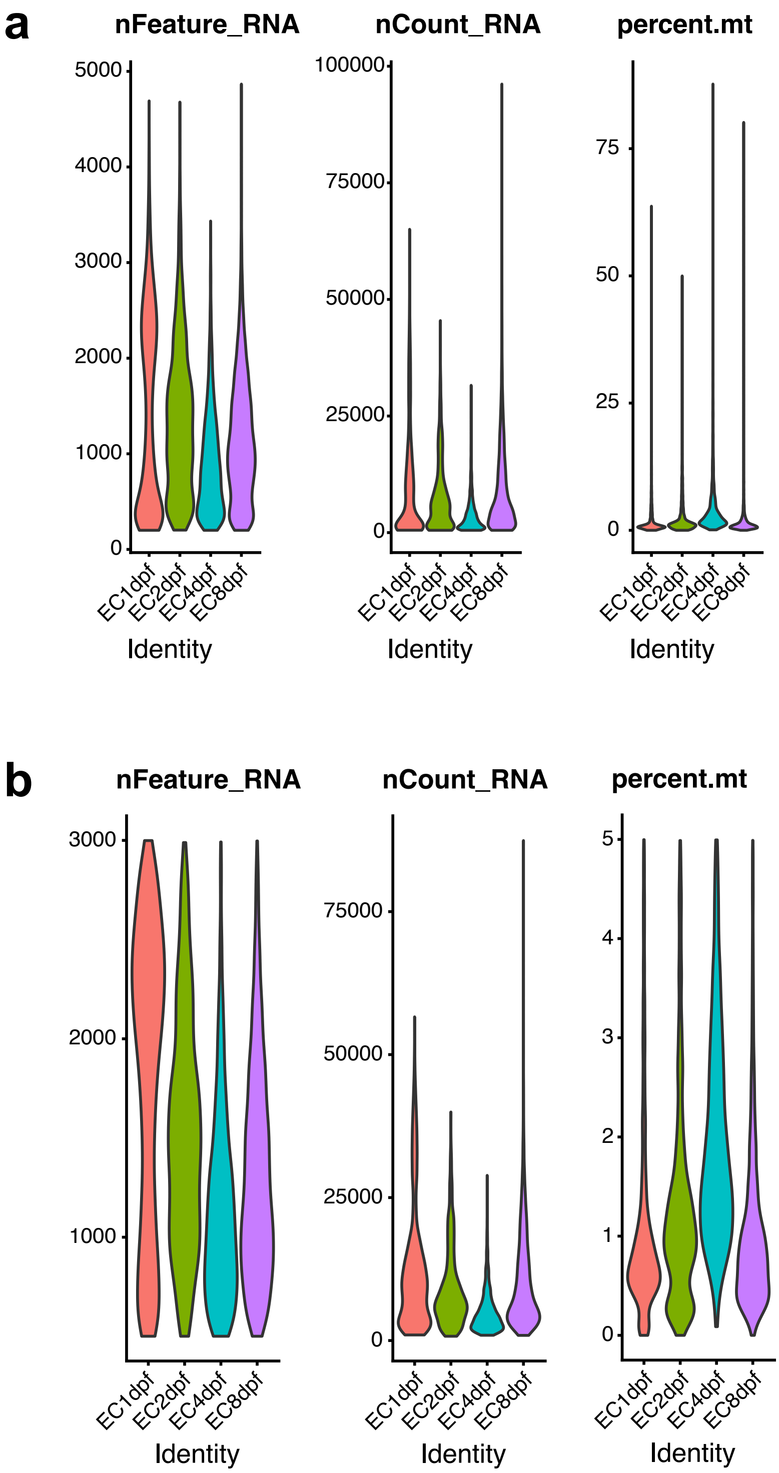

Supplementary Figure S3

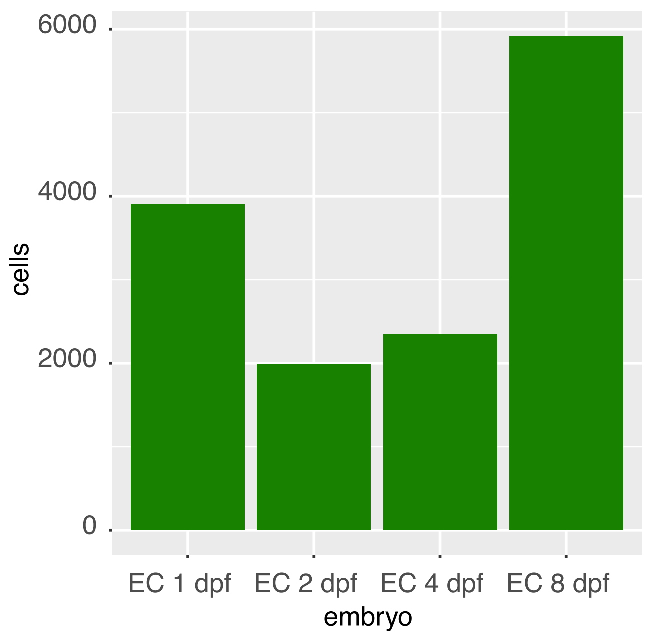

Supplementary Figure S4

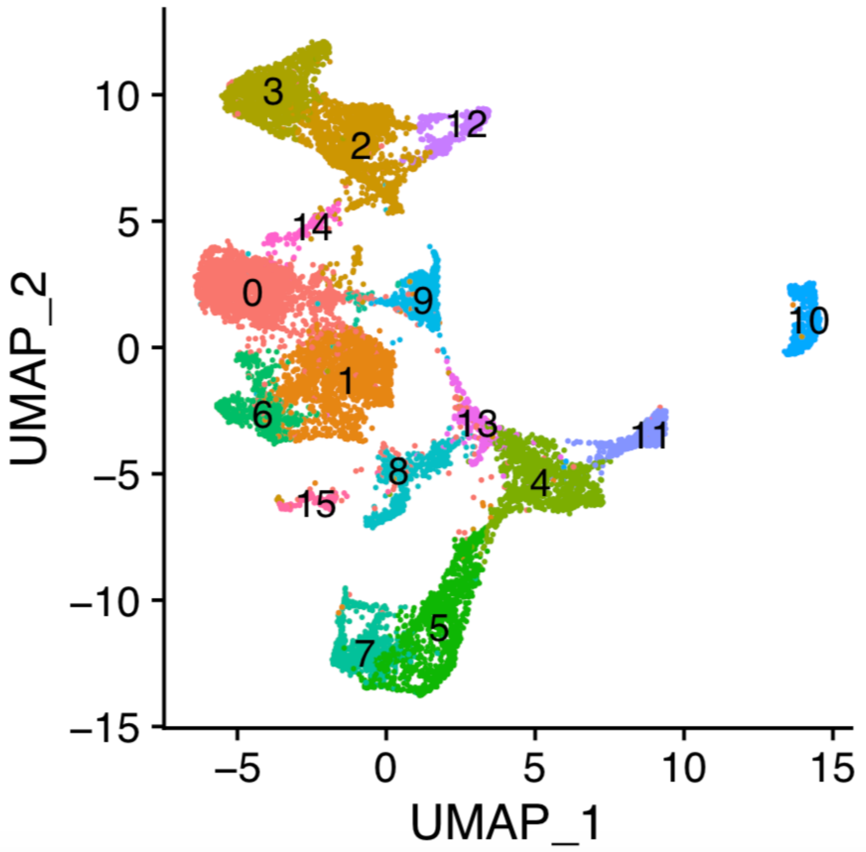


Total number of cells: 14168

Supplementary Figure S5

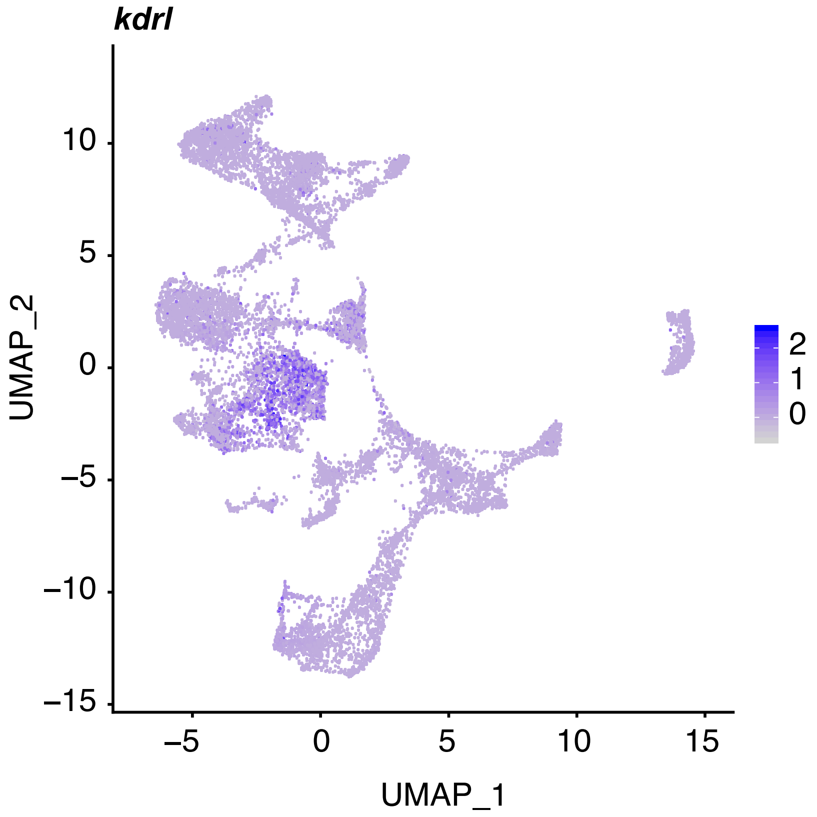

#### Supplementary Figure S6

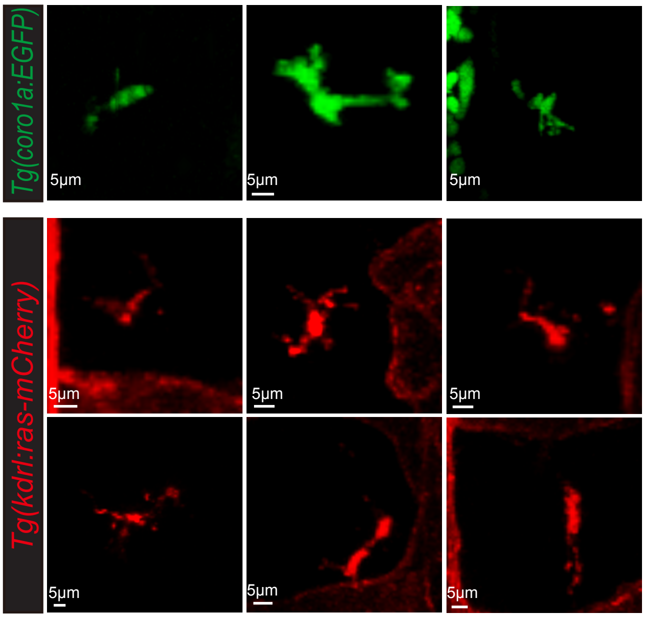

Supplementary Figure S7

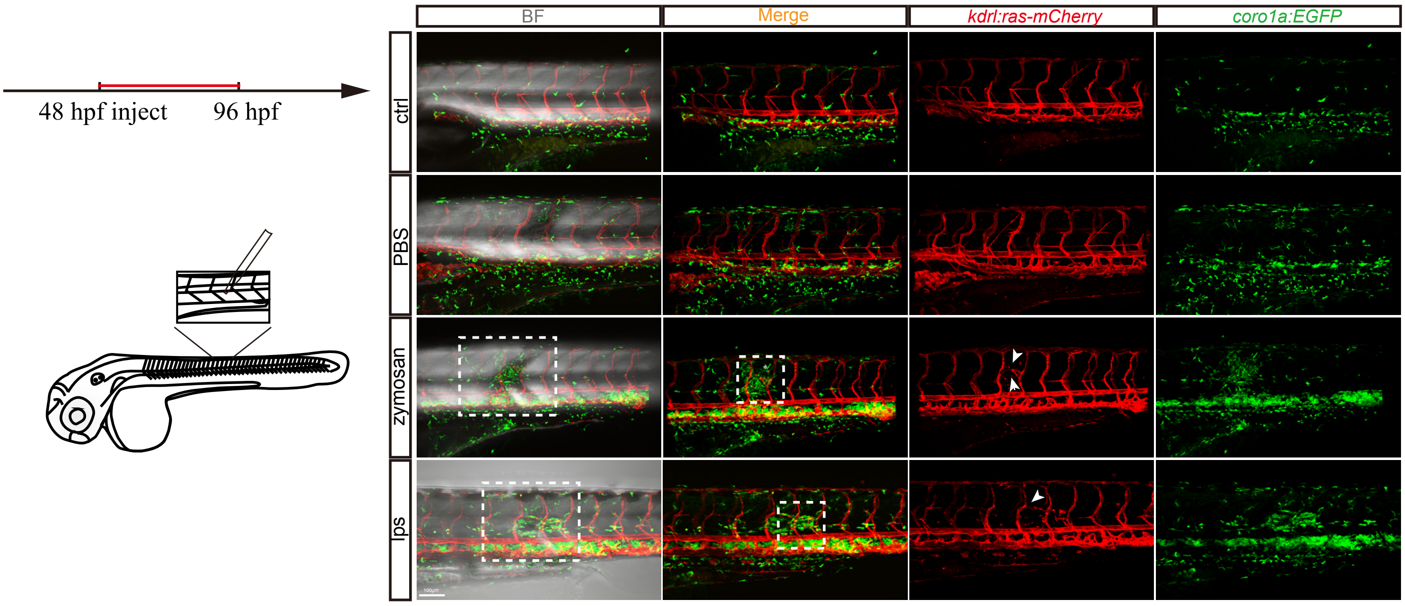
